# Transcriptional network analysis of PTEN protein-deficient prostate tumors reveals robust stromal reprogramming and signs of senescent paracrine communication

**DOI:** 10.1101/2025.01.31.635608

**Authors:** Ivana Rondon-Lorefice, Jose I. Lopez, Aitziber Ugalde-Olano, Maite Zufiaurre, Ianire Astobiza, Saioa Garcia-Longarte, Amaia Zabala, Sofia Rey, Aida Santos-Martin, Miguel Unda, Ana Loizaga-Iriarte, Mariona Graupera, Paolo Nuciforo, Arkaitz Carracedo, Isabel Mendizabal

## Abstract

Among the extensive genomic alterations in prostate cancer, the genomic deletion of *PTEN* stands out as one of the most consistently observed and confirmed alteration. *PTEN* loss in prostate tumors is primarily associated with cancer cell proliferation and survival through the activation of the PI3K-AKT-mTOR signalling pathway. However, its use as a robust biomarker in the clinical practice is hampered by the complex epigenetic, transcriptional and post-translational regulation. Here, we undertook an approach that combined *in situ* assessment of PTEN protein with transcriptional surrogates of its activity to gain insights into the downstream functional effects of PTEN loss in primary tumors. Our extensive bioinformatic analyses, including the integration with single-cell RNA-Seq approaches in a new clinical cohort, highlighted stroma remodeling as the major cancer cell-extrinsic process associated with PTEN loss. By applying similar computational strategy on the transcriptomic data generated from primary prostate tumors of genetically engineered *Pten knock-out* mouse models, we validated the causal role of *Pten* in the stromal reaction observed in clinical specimens. Mechanistically, we provide evidence for the activation of a paracrine program that encompasses enhanced TGF-β signalling and that is compatible with the secretome of PTEN-deficient senescent cancer cells. Our study provides relevant biological context to the cellular and molecular alterations unleashed upon PTEN protein loss.

## Introduction

Phosphatase and tensin homolog deleted in chromosome ten (*PTEN*) is a well-established tumor suppressor gene frequently disrupted in cancer. Germline and somatic mutations in *PTEN* have been associated with an increased risk for various types of malignancies. In prostate cancer, *PTEN* is one the most frequently altered genes in the epithelial gland, with 15-20% of patients presenting somatic genetic aberrations in primary tumors and up to 50% in patients with advanced disease (Hieronymus et al., 2017; Jamaspishvili et al., 2018; Taylor et al., 2010).

Loss of PTEN function results in the activation of the oncogenic PI3K signaling pathway. In prostate cancer, PTEN loss is associated to alterations in cell proliferation, migration, survival, metabolism, and genomic stability, among others (Lee, Chen and Pandolfi, 2018a). In addition to these well-characterized cancer cell-autonomous processes, recent studies have established critical roles of PTEN in the reprogramming of the tumor microenvironment, promoting immunosuppressive and pro-tumorigenic stroma in the prostate (Jamaspishvili, David M. Berman, *et al*., 2018; Vidotto *et al*., 2019).

The characterization of PTEN status in prostate cancer predominantly focuses on genomic loss. In multiple clinical cohorts, deletion of PTEN is strongly associated with a higher Gleason score (Wang and Dai, 2015) and poorer prognosis (Lotan *et al*., 2016; Bazzichetto *et al*., 2019). Yet, the assessment of *PTEN* gene status has not been translated into a robust prognostic or therapeutic biomarker in the clinical practice. This is possibly due to the highly complex regulation of its functions. *PTEN* regulation by promoter DNA methylation and non-coding RNAs such as miR-106a (Lu *et al*., 2019), miR-26a (Tian *et al*., 2013), and PlncRNA-1 (Cui *et al*., 2021) or competitive endogenous RNAs (Poliseno *et al*., 2010) have been reported, as well as several post-translational modifications including phosphorylation (Vazquez *et al*., 2000) and ubiquitination (Huang *et al*., 2012). Hence, genomic status might not fully capture the molecular and pathological consequences of loss of *PTEN* activity in prostate cancer.

Assessing PTEN protein levels in tissues can be more relevant as a clinical marker to develop new strategies for diagnosis and treatment. For example, PTEN protein status could guide stratification in clinical trials for treatment drugs such as PI3K or mTOR inhibitors. In addition, previous studies have shown that PTEN loss is frequently focal and subclonal in prostate tumors (Lotan *et al*., 2011; Gumuskaya *et al*., 2013). In this context, *in situ* methodologies such as immunohistochemistry (IHC) can be preferable to spatial-agnostic assays.

To obtain scientifically robust information around the molecular consequences of PTEN loss on prostate cancer biology, we combined IHC assays of PTEN with next-generation RNA sequencing technologies in a clinical cohort of 198 localized prostate cancer patients. By applying computational analyses and integrating single-cell transcriptomics, we identified stroma remodeling as a predominant consequence of PTEN protein loss in prostate tumors. We further capitalized on genetically engineered mouse models to infer the causal contribution of *Pten* loss to prostate tumor stroma remodeling.

## Results

### PTEN protein is extensively lost in prostate primary tumors

To assess the loss of PTEN protein in prostate tumors, we assembled a tissue microarray from a cohort of 198 patients from Basurto University Hospital with newly diagnosed disease (**Table 1** and **Supplementary Table 1** for detailed clinical and pathological characteristics of the cohort). We performed IHC in formalin-fixed and paraffin-embedded (FFPE) blocks obtained from prostatectomies. For most patients, two cylindrical sections (cores) per individual were available. The pathologist evaluated each core and assessed PTEN immunoreactivity. Using hematoxylin counterstaining, pathologists localized tumoral areas and evaluated PTEN staining in stromal regions as proxy for proper technical performance.

Out of 198 samples, 147 presented at least one tumor core with positive stromal PTEN staining (**Figure 1A**). Additional 51 samples were discarded due to tumor absence or negative stromal PTEN immunoreactivity (**Figure 1A, B** and **Supplementary Table 2**). For tumoral areas with positive stroma staining, the percentage of epithelial cells showing negative, weak, moderate, and intense PTEN staining was used to calculate the H-score (**Figure 1C**). Immunostaining values for PTEN in the primary prostate tumors showed a skewed distribution towards lower H-scores (median=5, mean 32.69, s.d.=50.86, **Figure 1D** left and **Supplementary Table 1**).

**Figure 1.**
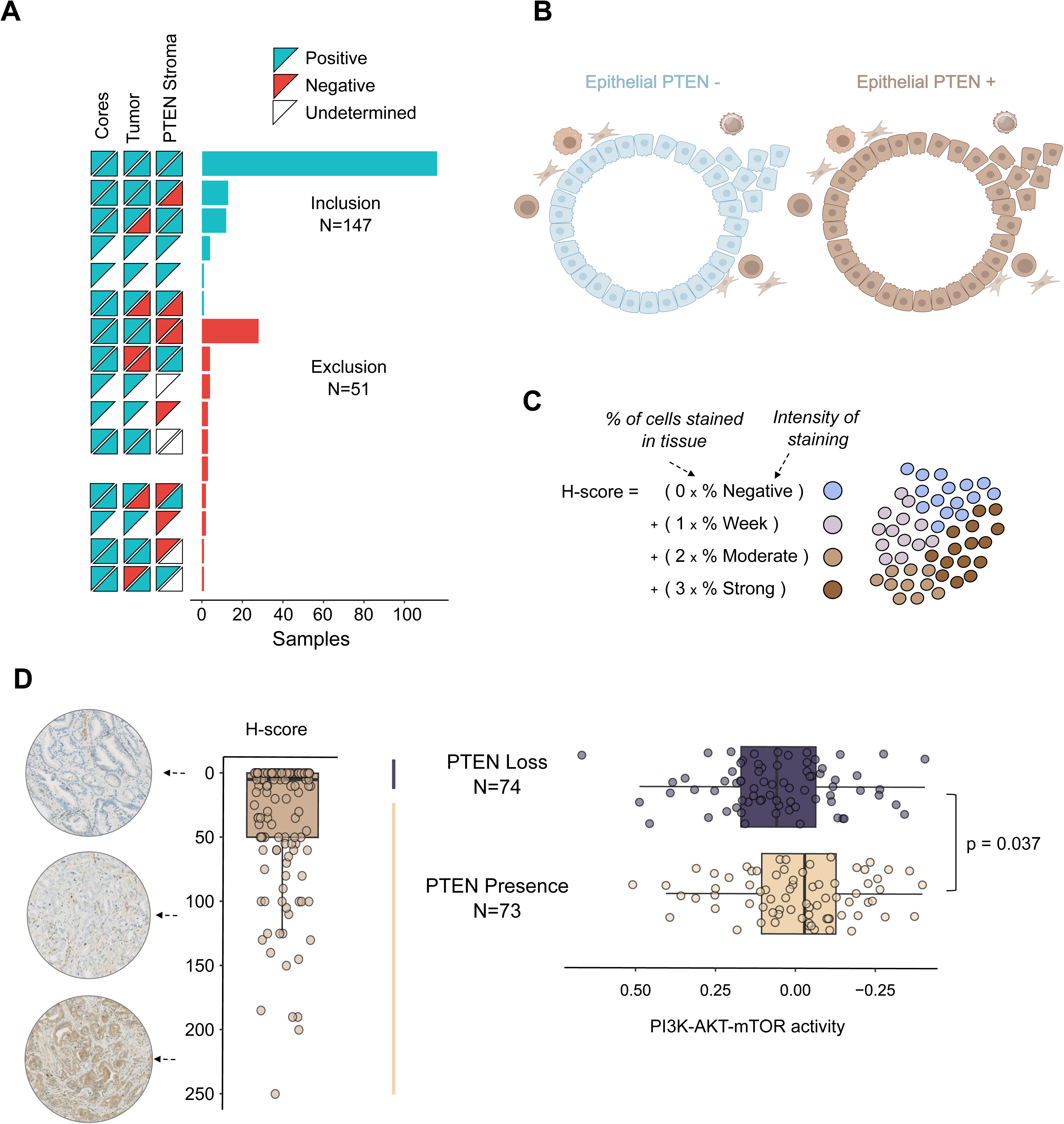
PTEN protein loss assessment in clinical prostate cancer specimens. **A.** Tissue microarrays from prostate specimens obtained from prostatectomies were employed to assess the status of PTEN by immunohistochemistry. Samples that failed to present at least one core with proper counterstaining, presence of tumoral areas, or showed negative staining for PTEN in the stromal cells were discarded from downstream analysis. **B.** Graphical representation of the stromal staining as quality control for PTEN signal presence (PTEN +) or loss (PTEN -) in the prostate epithelia. **C.** The H-score was computed to measure the staining of each sample based on the number of cells stained in the tissue and their intensity. **D**. Distribution of H-scores across the 147 prostate cancer samples. Arrows indicate representative images from the IHC assay. High H-scores correspond to strong PTEN staining (brown), while low H-scores indicate the absence of PTEN protein (blue). The cut threshold at H-score of 0 was employed to categorize the samples into PTEN loss and PTEN presence groups, that showed significant differences in PI3K-AKT-mTOR signature (p-value from two-sided Wilcoxon’s t-test).

### PTEN protein loss associates with extracellular-matrix processes

To understand the impact of PTEN protein loss on the transcriptomic landscape of prostate tumors we combined the immunoreactivity of the protein with RNA sequencing on this same cohort. Considering the canonical activation of PI3K-mTOR pathway upon PTEN loss (Lee, Chen and Pandolfi, 2018b), we leveraged on a PI3K-AKT-mTOR activity signature comprised of 105 genes (Liberzon *et al*., 2015) and obtained a binary classification of PTEN immunoreactivity (i.e. PTEN protein loss vs. PTEN protein presence). We monitored the statistical differences in the mean expression of PI3K-AKT-mTOR activity between PTEN loss and PTEN presence groups at various thresholds of the H-score (**Supplementary Figure 1A**). We identified that an H-score of 0 showed the best discriminative power, roughly distributing half of the samples per category (**Figure 1C** and **Supplementary Table 1**). Of note, *PTEN* mRNA levels did not vary according to PTEN protein status (**Supplementary Figure 1B**). These results suggest that PTEN protein detection by IHC is a good proxy of PI3K-AKT-mTOR activity while *PTEN* mRNA levels may not fully capture the complexity of post-transcriptional regulation.

To obtain a pathway-centered view of differential expression patterns associated with PTEN loss, we run Gene Set Enrichment Analyses (GSEA). The most significantly enriched pathways were associated with well-established functions of PTEN in the epithelia (such as “*Epithelial-mesenchymal tran*sition” **Figure 2A**). Interestingly, stromal-related functions were also highly enriched, including extracellular matrix (ECM)-related processes (**Figure 2A** and **Supplementary Table 3**). We next employed an unsupervised network-based approach without pre-defined gene categories or pathways. Weighted Gene Correlation Network Analyses WGCNA (Langfelder and Horvath, 2008) defines groups of genes (“*modules*”) characterized by similar expression patterns that potentially participate in specific biological pathways in a coordinated fashion. Out of the 22 modules identified, the modules labelled “*purple”* and “*green”*, exhibited the highest association with PTEN protein loss (Pearson’s r > 0.20, FDR < 0.05, **Figure 2B**, **Supplementary Figure 2**, and **Supplementary Tables 4, 5**).

**Figure 2.**
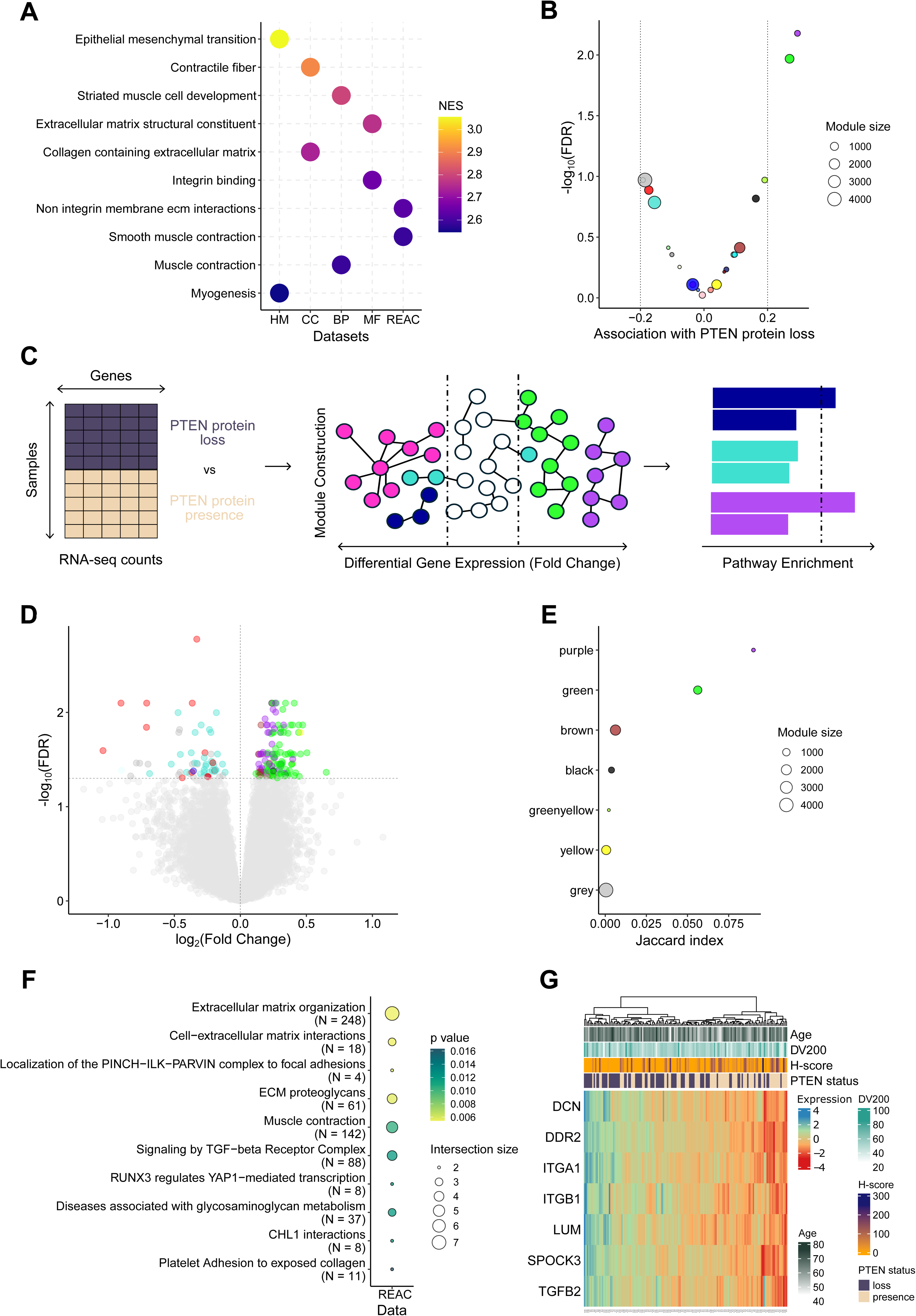
PTEN protein loss associates with extracellular matrix processes in prostate primary tumors. **A.** Gene-set Enrichment Analysis (GSEA) on gene expression data shows enrichment in extracellular matrix (ECM) related processes in PTEN protein loss versus presence. **B.** Volcano plot showing the results of the association (Pearson’s r) of each WGCNA module with the PTEN protein loss status. Dot size represents the number of genes in the modules. **C.** Volcano plot of differential gene expression analysis for PTEN loss vs PTEN intact status. The dots are colored according to their module membership of the genes in the WGCNA analyses. Lines represent statistical significance thresholds at Pearson’s r > 0.2 and FDR<0.05. **D**. Illustration of the workflow that combines co-expressed gene module identification with differential gene expression and downstream functional enrichment annotation. **E.** Intersection between modules and differentially expressed genes (DEGs) measured by Jaccard Index. Green and purple modules present the highest intersection with DEGs. **F**. Enrichment analysis conducted in the Reactome database shows overrepresentation of ECM related pathways. **G**. Heatmap illustrating the expression patterns of 7 differentially expressed genes in the green module enriched in the “*Extracellular matrix organization*” pathway.

We hypothesized that the genes showing significant differential expression at both the individual gene and network levels represent the subset with the highest potential to inform about the altered molecular landscape resulting from PTEN protein loss (**Figure 2C**). Gene-by-gene differential gene expression analyses (203 significant genes, |log_2_(fold change)|>0, FDR<0.05) showed that top overexpressed genes in tumors with PTEN protein loss were enriched in the purple and green upregulated modules (**Figure 2D, E**, **Supplementary Table 6)**. Functional enrichment of differentially expressed genes in the green module exhibited significant overrepresentation of extracellular matrix related functions (**Figure 2F, G**, and **Supplementary Table 7**). Instead, the purple module exhibited a notably higher enrichment in proliferative characteristics associated with cancer cells, such as the “*Activation of receptor-tyrosine kinases*” and “*PI3K signaling through fibroblast growth factor receptors*” (**Supplementary** Figure 3A, B). The information derived from the purple module is consistent with our threshold selection based on PI3K-AKT-mTOR activity signature and aligns with biological processes associated with PTEN loss described in the literature. Consequently, we chose to focus on the green module to explore the less understood roles of PTEN protein loss in the tumor microenvironment.

### PTEN protein loss associates with stromal remodeling

Given the functional link between ECM processes and the stromal compartment, we hypothesized that PTEN protein loss could have an impact in the relative abundance of the diverse cell types in the prostate tumor. To explore the compositional differences, we employed *in silico* deconvolution methods that infer cell-type proportions from bulk transcriptomic data. Tumor purity estimates (namely, the relative abundance of the epithelial compartment) (Yoshihara *et al*., 2013) were significantly decreased in the samples with PTEN protein loss, consistent with an increase in immune and stromal infiltration scores (**Figure 3A** and **Supplementary Table 8**). More detailed estimates for the abundances of 64 distinct cell-types with xCell (Aran, Hu and Butte, 2017) showed a notable increase in the relative abundance of fibroblasts in tumors exhibiting PTEN protein loss (**Figure 3B, Supplementary Tables 9, 10**). The deconvolution inference and ECM-related functions jointly suggest that the transcriptomic landscape of PTEN protein deficient tumors involves compositional alterations in the tumor microenvironment.

**Figure 3.**
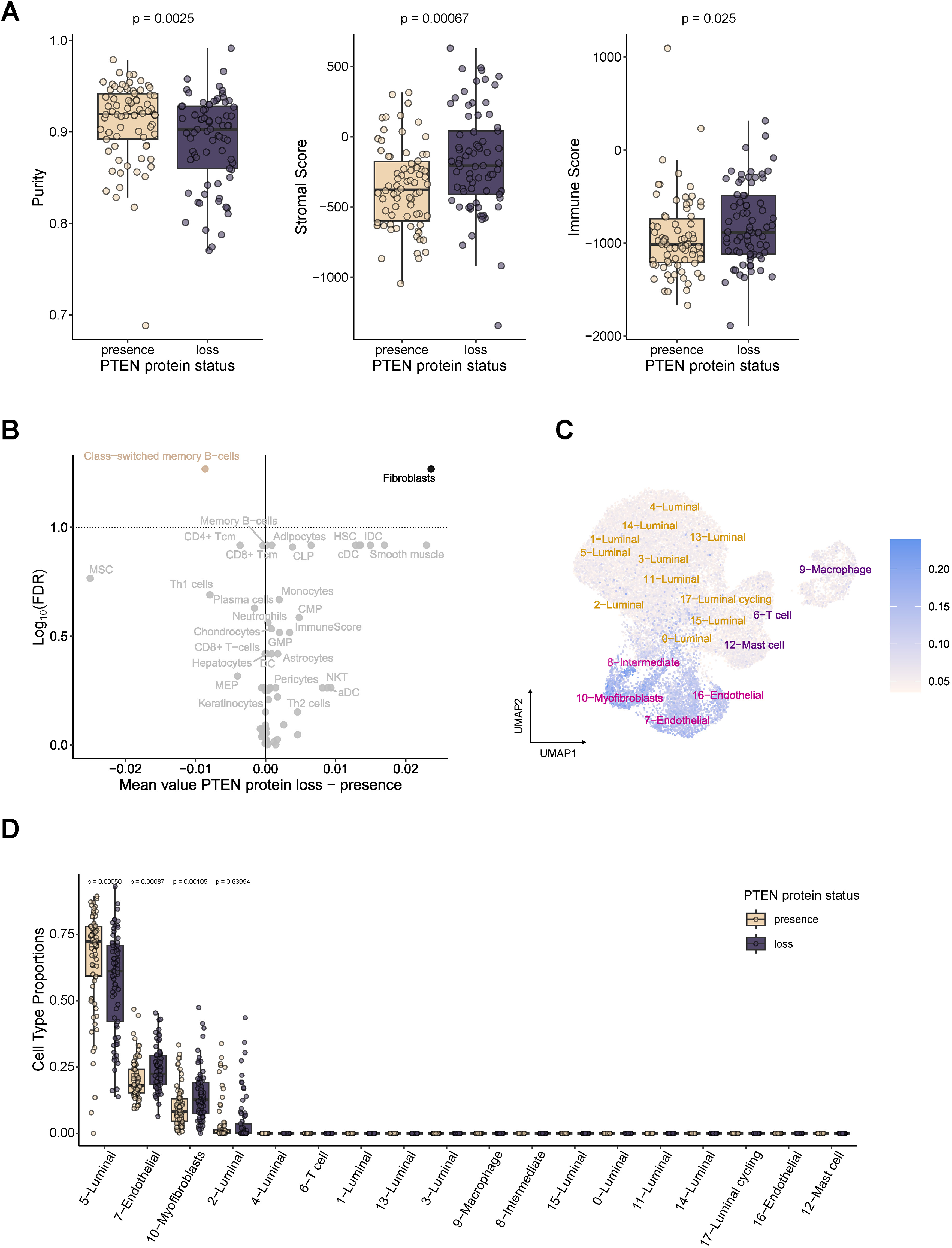
Increase in stromal composition observed with PTEN protein loss in prostate cancer tumors. **A.** Cellular decomposition analysis showing the purity (the percentage of epithelial cells in the tumor) and the estimates for stromal and immune abundances using ESTIMATE. **B.** Detailed decomposition into specific cell types shows a significant enrichment for fibroblasts at primary tumors without PTEN protein expression using xCell**. C.** UMAP projection of a single-cell RNA-Seq database comprising 8 primary prostate specimens. The plot shows the expression level of differentially expressed genes in the green module. **D.** Prostate-specific decomposition analysis performed on a single-cell dataset from 8 primary prostate specimens.

To refine these findings, we next leveraged on publicly available single-cell RNA-Seq data from prostate primary tumors (Chen *et al*., 2021) to perform cell-type deconvolution(**Supplementary Figure 3C-D)**. .Unlike xCell, which relies on bulk transcriptomic data and predefined marker genes, MuSiC incorporates single-cell expression profiles to improve cell-type resolution (Wang *et al*., 2019). Consistent with our previous results, we observed a decrease in the luminal compartment and an increase in the endothelial and myofibroblast compartments in PTEN protein deficient tumors (**Figure 3C**). Next, we analysed the expression breadth of the genes impacted by PTEN protein loss with single-cell resolution. Cells with the highest expression levels for the DEGs in the green module were confined to the non-immune stromal compartment (**Figure 3D**). Altogether, cell-deconvolution and single-cell resolution analyses support that PTEN protein loss in the prostate epithelium plays a major role in driving a dynamic response of the non-immune tumor microenvironment, influencing the abundance of cancer-associated fibroblasts and the tumor vasculature.

### PTEN loss induces stromal remodeling in mouse models

Our analyses of human transcriptomes unveiled an association between PTEN loss and stromal remodeling. However, considering the complex genomic landscape of prostate adenocarcinoma (Campbell *et al*., 2017) clinical data does not allow us to establish a direct causal relationship. To this end, we leveraged experimental murine models of prostate cancer with conditional *Pten* deletion specific to prostatic epithelial cells (*Pten^flox^*by Cre recombinase under the control of the Probasin promoter). Generally, Pten *^pc−/−^* mice develop prostate intraepithelial lesions (PIN) by the age of 3 months, with cancer manifesting by 6 months (**Figure 4A**) (Chen *et al*., 2005). As predicted, an increase of the PI3K-AKT-mTOR signaling (Liberzon *et al*., 2015) was also evident in the transcriptomes of Pten-deficient prostates at 6 months (**Figure 4A**).

**Figure 4.**
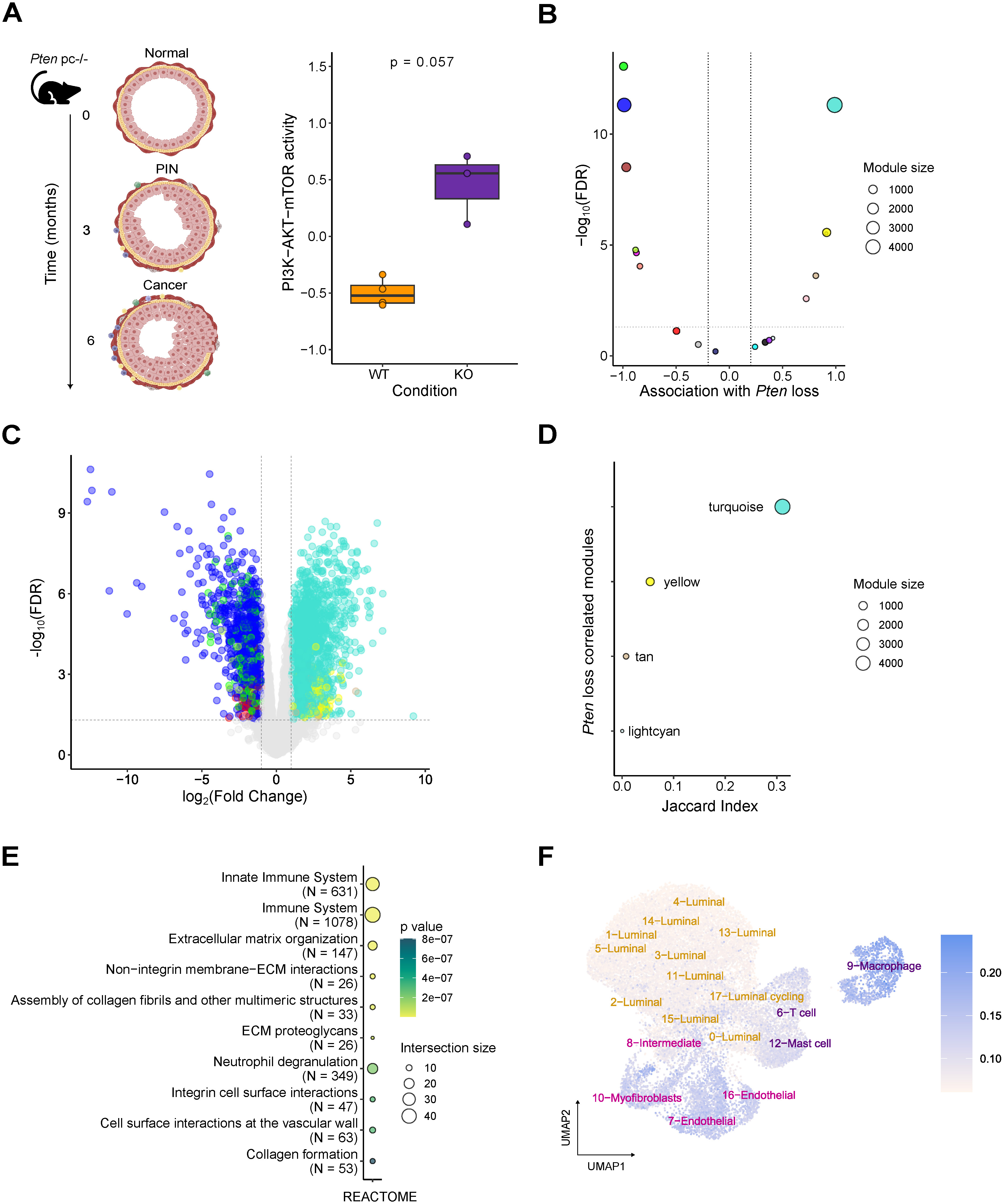
*Pten* loss drives stromal changes in mouse models. **A.** Left: Illustration of the progression of a conditional Pten mouse model for prostate cancer. Mouse models with homozygous deletion of *Pten* in the epithelial compartment typically develop prostate cancer by the age of 6 months. Right: *Pten knock-out* mice show increased PI3K-AKT-mTOR signaling in the prostate epithelia at 6 months (Wilcoxon two-tailed test). **B.** The association of WGCNA modules (Pearson’s r) with Pten null status. Lines represent statistical significance thresholds at Pearson’s r > 0.2 and FDR<0.05. **C.** Volcano plot of differential gene expression analysis for PTEN loss vs PTEN intact status. The dots are colored according to their module membership of the genes in the WGCNA analyses. Lines represent statistical significance thresholds |log_2_(fold change)| > 0 and FDR<0.05. **D**. Intersection analysis between modules and differentially expressed genes (DEGs) measured by the Jaccard Index. Turquoise and yellow modules present the highest proportion of DEGs. E. Module yellow exhibits significant enrichment in extracellular matrix (ECM)-related pathways, suggesting a potential role in stromal remodeling. **F**. UMAP projection of a single-cell RNA-Seq database comprising 8 primary prostate specimens. The plot shows the expression levels of differentially expressed genes within the yellow module.

We applied the module construction and differential gene expression workflow used in the human datasets (outlined in **Figure 2E**) to murine prostate transcriptomic profiles of from *Pten* knock-out and wild-type mice at 6 months. Two modules exhibited the strongest association with *Pten* loss in the murine model, which also presented the highest amount of differentially expressed genes (**Figure 4B-D, Supplementary Figure 4, Supplementary Tables 11-13**). The first module, turquoise, was notably large, comprising 4,680 genes and 1,749 DEGs, was primarily enriched in immune-related pathways (**Supplementary Figure 5**). Interestingly, the second module, labelled yellow, exhibited significant enrichment in ECM-associated mechanisms such as “*Extracellular matrix organization”* (**Figure 4E, Supplementary Table 14**). Upon converting the genes in this module to their human homologues, we found that these were predominantly expressed in macrophages, endothelial cells and myofibroblasts in single-cell data (**Figure 4F**). Further deconvolution analysis using single-cell RNA-Seq revealed increased stromal cell contribution in *Pten*-deficient mice (**Supplementary Figure 6**). These observations suggest that the absence of PTEN protein in the prostate epithelia induces critical changes in the stromal composition of tumors in both experimental models and clinical specimens.

### PTEN protein loss in tumors is associated with activated TGF-**β** senescence-associated secretory phenotype programs

To ascertain the molecular means behind the stroma remodeling induced upon tumor cell-intrinsic PTEN protein loss, we focused on paracrine signals that could explain this phenomenon. Altered secretome profiles have been reported in PTEN-deficient cells concomitant to the induction of cellular senescence(Lee, Chen and Pandolfi, 2018a). Senescence is a combination of cell-intrinsic arrest combined with a paracrine secretory program termed senescence-associated secretory phenotype (SASP) (Gorgoulis *et al*., 2019).Interestingly, upon analysis of the differential transcriptome of PTEN-deficient prostate cancer clinical specimens we found a remarkable enrichment in activated TGF-β (Transforming Growth Factor-β) pathway (**Figure 2E**). TGF-β participates in the senescence response and orchestrates the remodeling of the extracellular matrix and the stroma. Hence, we hypothesized that altered TGF-β transcriptional programs and stroma remodeling were surrogate signals of the secretory reprogramming of PTEN deficient tumors associated to the SASP. To this end, we explored the association of PTEN protein loss with senescence (Troiani *et al*., 2022) and TGF-β activation levels (Calon *et al*., 2012) in different pre-clinical and clinical transcriptomics datasets. In our patient cohort, we consistently observed an upregulation of these pathways in the primary tumors exhibiting PTEN protein loss and a strong correlation with the stromal remodeling signature (green module) (**Figure 5A, D, and Supplementary Figure 7**). Interestingly, the associations of stromal remodeling with senescence and TGF-β were validated in the TCGA PRAD cohort, yet the association with genetic loss was not significant (**Figure 5B, E and Supplementary Figure 7**). Consistently, we also observed these trends in our mouse models upon *Pten* deletion (**Figure 5C, F**). Altogether, these data suggest that PTEN protein loss is a robust indicator of the activation of senescence-associated processes in localized prostate cancer, and that the link between SASP and stroma remodeling emerges as one of the most prominent molecular programs in PTEN protein-deficient tumors.

**Figure 5.**
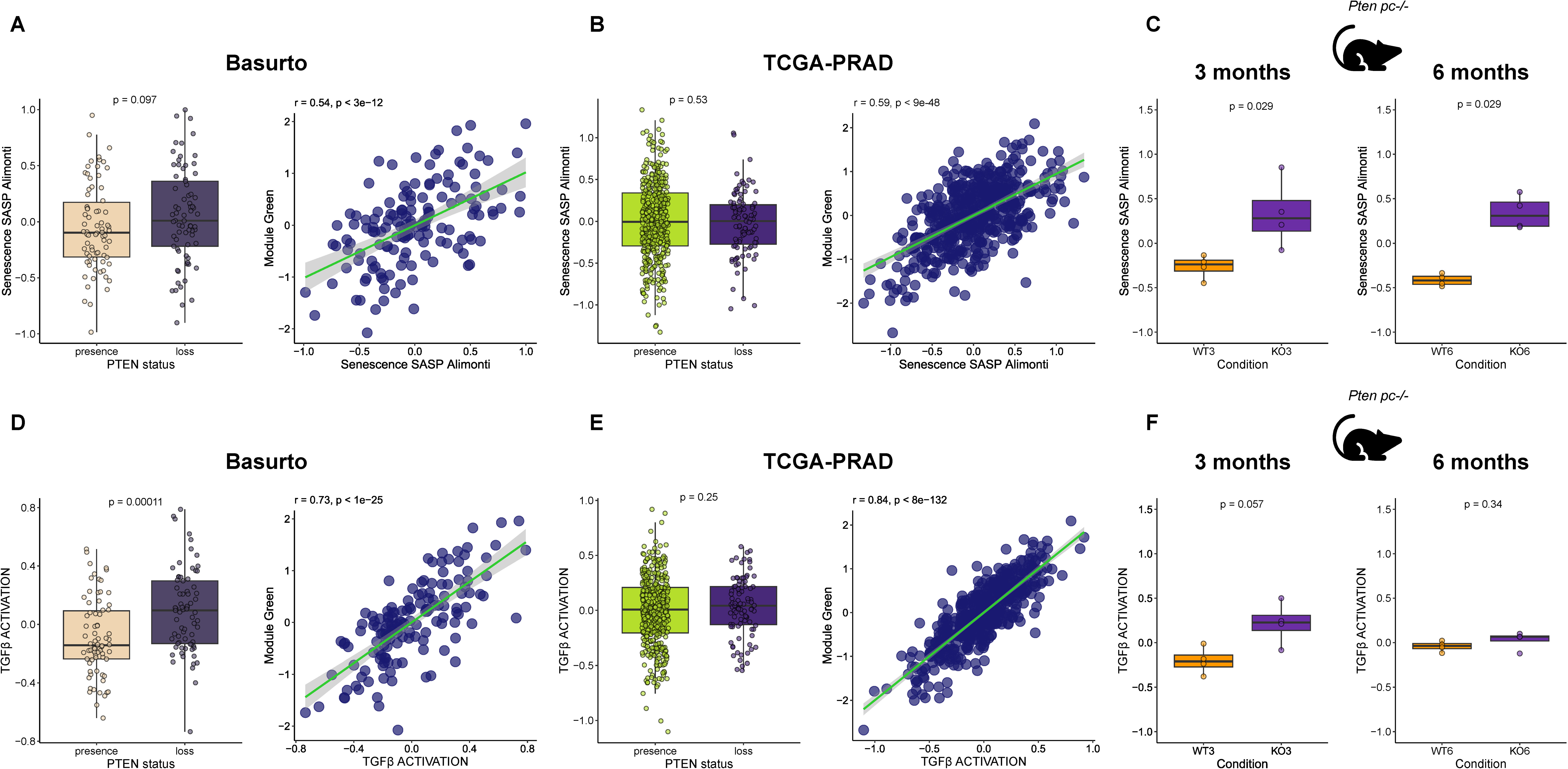
PTEN protein loss associates with senescence and TGF-β signatures. **A.** Left: Senescence-associated secretory phenotype (SASP) signature in our patient cohort according to PTEN protein loss and presence. Right: correlation between the SASP signature and the signature based on the DEGs in the green module in our cohort. **B.** Left: Senescence-associated secretory phenotype (SASP) signature in the TCGA patient cohort according to PTEN gene copy loss. Right: correlation between the SASP signature and the signature based on the DEGs in the green module in the TCGA. **C.** Left: Senescence-associated secretory phenotype (SASP) signature in the animal model with Pten deletion and wild type at three (left) and nine months (right). **D.** Left: TGF-β signature in our patient cohort according to PTEN protein loss and presence. Right: correlation between the TGF-β signature and the signature based on the DEGs in the green module in our cohort. **E.** Left: TGF-β signature in the TCGA-PRAD patient cohort according to PTEN gene copy loss. Right: correlation between the TGF-β signature and the signature based on the DEGs in the green module in the TCGA-PRAD cohort. **F.** Left: TGF-β signature in the animal model with Pten deletion and wild type at three (left) and nine months (right)

## Discussion

Genomic loss of *PTEN* represents one of the most robustly reproducible and extensively validated genetic alterations in prostate cancer. In this study we assembled a novel prostate cancer clinical cohort (n=198, from Basurto University Hospital in Spain) to quantify the loss of PTEN protein by immunohistochemistry in primary tumor specimens derived from radical prostatectomy. We adhered to stringent quality criteria to ensure maximum robustness of our assessment of PTEN protein loss. First, we followed a clinically approved PTEN IHC protocol from Vall d’Hebron Hospital in Barcelona, Spain (Sauri *et al*., 2016). Second, we utilized stromal PTEN positive staining as a quality control of the integrity and preservation of the tissue prior to epithelial assessment of the phosphatase. Third, we coupled the IHC analysis with RNA-Seq analysis of the same specimens to perform a potent computational integrative study. These quality standards are critical to making reliable assessments of PTEN protein loss. On the one hand, within primary prostate tumors, different areas often exhibit varying levels of *PTEN* gene loss. Remarkably, intratumoral heterogeneity in genomic loss of PTEN has been reported in half of the cases (Gumuskaya *et al*., 2013; Krohn *et al*., 2014; Ahearn *et al*., 2016). The inclusion of two tumor cores along with the criteria of positive staining of the stroma across different tumoral regions as a control, can mitigate these potential limitations in our study. On the other hand, IHC may fail to detect post-translational modifications that can influence PTEN protein activity, stability, and half-life. By integrating the H-scores with a surrogate marker of PI3K-AKT-mTOR pathway transcriptional activity, we achieved a binary classification of PTEN protein loss that also accounts for the activation state of major effector pathways.

Our analyses identified PTEN protein loss in about 50% of the patients in our cohort. This PTEN loss at the protein level is higher than the loss reported at the genomic level in primary tumors (15-20% (Taylor *et al*., 2010; Hieronymus *et al*., 2017; Jamaspishvili, David M Berman, *et al*., 2018)) while in line with previous reports. For example, Lotan et al. (Lotan *et al*., 2017) reported that only 66% of primary tumors with PTEN protein loss presented *PTEN* genomic deletion by fluorescence *in situ* hybridization. This discrepancy supports the need of tissue-based protein assays to capture the complexities of PTEN activity loss within prostate primary tumors.

We further capitalized in our RNA-Seq data to identify transcriptome-wide changes associated with PTEN protein loss. By combining pathway-based and agnostic gene-network approaches, we identified a group of co-expressed genes related to extracellular matrix functions (ECM) as the main cancer cell-extrinsic process differentially associated with PTEN protein loss. *In silico* cell-type deconvolution analyses and integration with transcriptomic datasets at single-cell resolution, revealed an expansion of the stromal compartment in association with PTEN protein loss and a concomitant upregulation of ECM-associated genes that was confined to the non-immune stroma.

Prostate primary tumors harbor numerous genomic, transcriptomic and proteomic alterations, with PTEN loss co-occurring alongside these changes. This complexity highlights the correlative nature of clinical studies, which we mitigated through the implementation of animal models with tissue specific deletion of *Pten*, thus allowing us to extract those molecular process in human prostate cancer that are causally associated to PTEN loss. By employing genetically modified prostate-specific *PTEN* knockout mice, we observed a tumoral transcriptomic rewiring and stromal reprogramming that matched our observations in human specimens. Together, these data suggest a causal link between PTEN loss and stromal reaction in prostate cancer. These patterns agree with the phenomenon of stromal reaction described in several solid tumors including the prostate, linked to tumor growth, invasion, and metastasis (Quail and Joyce, 2013) (Tuxhorn, Ayala, *et al*., 2002). Reactive stroma in prostate cancer involves a transition of cancer-associated fibroblasts (CAFs) into myofibroblast and inflammatory CAFs, remodelled ECM, activated angiogenic niche and an immunosuppressive landscape (Tuxhorn, Ayala, *et al*., 2002; Tuxhorn, McAlhany, *et al*., 2002).

By presenting a publicly accessible, novel patient cohort with integrated IHC and RNA-Seq data, our study comprehensively explores the contribution of PTEN protein loss to key processes in prostate cancer. Mechanistically, we identified significant enrichment of the TGF-β signaling pathway in an ECM-like gene program upregulated upon PTEN loss. The positive correlation between TGF-β activity, senescence signatures, and PTEN loss in human cohorts and murine models suggested that PTEN loss in epithelial cells promotes a paracrine program potentially emanating from senescent tumor cells that governs the remodeling of the stroma. We propose that this reaction is mediated by the activation of TGF-β signaling within the SASP, which enhances the stromal reaction and ECM remodeling in the tumor microenvironment.

## Methods

### Tissue Collection

We collected 198 tissue specimens from localized primary prostate tumors from Basurto University Hospital in Bilbao (Spain). The tissues were derived from prostatectomies following (Ugalde-Olano *et al*., 2015) and were preserved in formalin-fixed paraffin-embedded (FFPE) blocks. Sample collection was coordinated by the Basque Biobank. The sample acquisition was conducted in accordance with the ethical guidelines and approved protocols CEIC-E 11-12, 14-14, and 19-20. See **Table 1** for detailed information of the clinical and pathological characteristics of the cohort. Tumor-rich regions were selected for RNA extraction by the pathologist.

### Immunohistochemical staining of PTEN

The full cohort was represented in ten tissue microarrays (TMA), in which each patient was represented with 2 tumor cell-rich cores of 1mm of diameter. The TMA slides were dewaxed at 72°C, incubated with an anti-PTEN rabbit monoclonal primary antibody (clone 138G6, 1/100 diluted, #9559 Cell Signaling Technology) for 60 min at 36 °C and detected using Ultraview DAB IHC detection Kit (Cat. No. 760-500) on the VENTANA BenchMark ULTRA automated staining platform. The slides were mounted and digitalized at 20x using a slide scanner (NanoZoomer 2.OHT, Hamamatsu Photonics, Japan).

### PTEN immunohistochemical scoring

Following digitalization, prostate tissues were assessed by a board-certified pathologist at the Cruces Hospital at a microscope. As technical controls, we evaluated the positive counterstain with haematoxylin, presence of tumoral regions, and PTEN positive staining in the stroma per core. A total of 51 samples did not pass these criteria in at least one core and were excluded for downstream analyses (**Figure 1A** and **Supplementary Table 2**).

With the 147 samples that passed the quality controls, the percentage of cells exhibiting negative, weak, moderate, and intense staining in the epithelia was then utilized to calculate the H-score, calculated as follows (where x means multiplication):

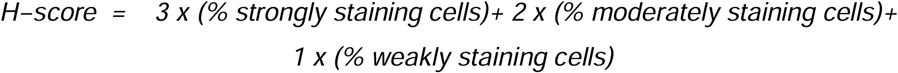

When two cores are available per patient the mean H-score is reported.

### RNA-Seq library generation and data processing of human specimens

RNA was extracted from paraffin-embedded tissue sections using a standardized protocol. Following deparaffinization, enzymatic digestion and heat treatments were applied to facilitate tissue breakdown and nucleic acid release. The RNA was purified using silica-based column technology, incorporating DNase treatment to remove contaminating DNA. Eluted RNA was assessed for integrity and concentration using spectrophotometry, gel electrophoresis, and capillary electrophoresis systems. Samples were stored at −80°C. Sequencing libraries were prepared following the SMARTer Stranded Total RNA-Seq Kit v2 - Pico Input Mammalian protocol (Takara BioUSA, Cat. #634411), as detailed in the “SMARTer Stranded Total RNA-Seq Kit v2 – Pico Input Mammalian User Manual (Rev. 050619). The librariess were sequenced using the NovaSeq 6000 platform with pared end reads of 150 base pairs in length (average 63 million reads per sample).

Adaptor sequences specified by the Illumina TruSeq RNA kit were removed using cutadapt (Martin, 2011) (v4.8) with the following command: cutadapt - a AGATCGGAAGAGCACACGTCTGAACTCCAGTCA-A AGATCGGAAGAGCGTCGTGTAGGGAAAGAGTGT. Additionally, three nucleotides were trimmed from the R1 and R2 strands, respectively, as specified by the library preparation protocol using the command: cutadapt -q 10 --minimum-length 20 -u-3 -U 3. The quality of the reads was verified using FastQC (v0.12.1).

Reads were mapped to the reference human genome GRCh38.v94 using the Spliced Transcripts Alignment to a Reference (STAR) (Dobin *et al*., 2013) 2.7.10 a with the following parameters: --outFilterMultimapNmax 1 --outReadsUnmapped Fastx --outSAMtype BAM SortedByCoordinate --twopassMode Basic --limitBAMsortRAM 2000000000 --quantMode TranscriptomeSAM GeneCounts.

### RNA-Seq library generation and data processing of murine data

RNA was extracted from (fresh) tissues and sequenced using the Illumina HiSeq4000 platform with paired-end reads of 100 base pairs in length (average 66 million reads per sample). Sequencing libraries were prepared following the “TruSeq Stranded Total RNA Human/Mouse/Rat” kit (Illumina Inc., Cat. #RS-122-2201), using “TruSeq Stranded Total RNA Sample Prep-guide (Part#15031048Rev.E)”. The quality of the reads was verified using FastQC (v0.12.1).

Reads were mapped to the reference mouse genome GRCm39 using the Spliced Transcripts Alignment to a Reference (STAR) (Dobin *et al*., 2013) 2.7.10 a with the following parameters: --outFilterMultimapNmax 1 --outReadsUnmapped Fastx --outSAMtype BAM SortedByCoordinate --twopassMode Basic --limitBAMsortRAM 2000000000 --quantMode TranscriptomeSAM GeneCounts.

### Differential expression and weighted-gene correlation network analyses (WGCNA)

We used limma voom from edgeR (Robinson, McCarthy and Smyth, 2010) (v4.0.16) to identify the differentially expressed genes (DEGs) between individuals with PTEN protein loss vs presence condition. In the case of the human dataset, we considered age and integrity index DV200 index (percentage of RNA fragments > 200 nucleotides (Matsubara *et al*., 2020)) as covariates in the analyses after Z-score transformation. Differentially expressed genes were identified using a threshold of |FC|>1 and FDR<0.05; and |FC|>2 and FDR<0.05 for human and mouse datasets, respectively.

We performed gene co-expression network analysis using WGCNA (Langfelder and Horvath, 2008) (V 1.72.1) in R. Briefly, in WGCNA, the construction of the network is based on soft thresholding the correlation coefficient to which the co-expression similarity is raised to calculate the adjacency. In the case of human data, we filtered out genes with less than 5 counts in more than 70% of the samples; for the mouse data, we filtered out genes with less than 5 counts in more than 90% of the samples. We subsequently applied variance-stabilization transformation to the gene expression matrices for both human and mouse using DESeq2 (Love, Huber and Anders, 2014) (v1.42.1).

To select the correct parameter for network construction, we inferred the soft-threshold power by approximating our network to a scale-free topology using a signed Spearman correlation. For the human dataset we selected a power of 14 based on the results obtained with the function ScaleFreeTopology. Then, we run the function blockwiseModule passing through the following parameters maxBlockSize= 25000, corType= “bicor”, maxPOutliers =0.1, pearsonFallback = “individual”, networkType =”signed”, deepSplit=1, mergeCutHeight = 0.1. In the mouse dataset the parameters passed to blockwiseModule function were the following: maxBlockSize=20000, corType = “bicor”, maxPOutliers = 0.1, power = 18, networkType =“signed”, deepSplit = 2, mergeCutHeight = 0.25. To intersect human and mouse modules, conversion of mouse gene names to human was performed using the gene homology data from Ensembl.

### Functional enrichment analyses

We used Gene Set Enrichment Analysis (GSEA) (Subramanian *et al*., 2005)(v 4.3.2) after applying the normalization method “median of ratios” with DESeq2 (Love, Huber and Anders, 2014) (v1.42.1). We identified the enriched pathways in PTEN protein loss vs presence in the following gene sets databases for Human Collection (MSigDB): h.all.v2023, c5.go.bp.v2023, c5.go.cc.v2023, c5.go.mf.v2023, c2.cp.reactome.v2023. We used the permutation type by “gene_set”.

Overrepresentation analyses were carried out with “gprofiler” package (Kolberg *et al*., 2023)(v0.7.0). For module enrichment analysis, the background was tuned by all genes evaluated in the module construction. The following parameters were selected: organism = “hsapiens” or “mmusculus”, multi_query = FALSE, significant = TRUE, exclude_iea = FALSE, measure_underrepresentation = FALSE, evcodes = TRUE, user_threshold = 0.05, correction_method = “fdr”, domain_scope = “custom”, numeric_ns = “”, sources = NULL, as_short_link = FALSE.

### In-silico deconvolution analyses

We estimated the purity (cancer cell proportion) per sample by applying tidyestimate (V 1.1.0) based on the gene-set-enrichment-analysis (GSEA) of stromal and immune gene sets. For an extensive cell-type deconvolution across 64 cell types, we ran xCell (Aran, Hu and Butte, 2017) (V 1.1.0) using transcripts per million (TPM) normalization. We further refined our analysis by estimating cell type proportions specific to prostate tissue using Music (V 1.0.0).

## Data and script availability

The data generated in this project (raw and processed) are available in SRA and GEO at accession numbers XXXX. The code for data analyses are available at https://github.com/imendizabalCIC/PTEN_protein_loss_project/.

## Authorś contributions

I. R.-L. and S.G.-L. performed bioinformatic analyses on bulk and single-cell transcriptomic datasets and contributed to the preparation of the figures for the manuscript. I.M. supervised the work of I. R.-L. and S.G.-L. M.U. and A.L.-I. generated the Basurto cohort and provided biological specimens supported by S.R. and A.S.-M. I.A. and A.Z. coordinated the experimental preparation of the IHC under the supervision of A.C. M.Z. and A.U-O prepared the TMAs. IHC experiments were performed by P.N. J.I:L. performed the IHC quantification. M.G. contributed to the study design. A.C. and I.M. conceived the study, supervised the execution of the project and wrote the manuscript. All authors have read and approved the final version of the manuscript.

---

**Table 1.** Clinical information of the patient cohort described in this study.

## Supplementary Figures

**Supplementary Figure 1. A.** The plot shows the p-value results for the Wilcoxon test between PTEN loss and PTEN presence for the PI3K-AKT-mTOR activity signature (y-axis) according to different H-score thresholds (x-axis). An H-score cut-off value of 0 displayed the largest differences between PTEN loss and PTEN presence categories. **B.** mRNA levels of *PTEN* do not show significant differences according to PTEN protein status (Wilcoxon test).

**Supplementary Figure 2. A.** Analysis of network topology for different soft-thresholding powers. The left panel shows the scale-free fit index (y-axis) as a function of the soft-thresholding power (x-axis). The right panel shows the mean connectivity (y-axis) as a function of the soft-thresholding power (x-axis). **B.** Clustering dendrogram of all genes based on the hierarchical clustering of adjacency-based dissimilarity. The coloured row below the dendrogram indicates module membership. **C.** Barplot showing the size (number of genes) of each module. **D.** Heatmap of correlations of eigengenes (first principal component) between the modules. **E.** WGCNA modules and their association with biological and clinical variables.

**Supplementary Figure 3. A**. Module purple is enriched in pathways related to Receptor Tyrosine Kinases signaling and PI3K activation by Fibroblast receptor factors. **B**. Heatmap of top differentially expressed genes in purple module. **C**. Dotplot of the markers used to annotate each cluster in single-cell data. **D**. Genes in the purple module are expressed mostly in non-immune and immune stroma.

**Supplementary Figure 4. A.** Analysis of network topology for different soft-thresholding powers. The left panel shows the scale-free fit index (y-axis) as a function of the soft-thresholding power (x-axis). The right panel shows the mean connectivity (y-axis) as a function of the soft-thresholding power (x-axis). **B.** Clustering dendrogram of all genes based on the hierarchical clustering of adjacency-based dissimilarity. The coloured row below the dendrogram indicates module membership. **C.** Barplot showing the size (number of genes) of each module. **D.** Heatmap of correlations of eigengenes (first principal component) between the modules. **E.** WGCNA modules and their association with biological and clinical variables.

**Supplementary Figure 5.** Functional enrichment analyses of the differentially expressed genes in the turquoise module.

**Supplementary Figure 6.** Deconvolution analysis of the mouse transcriptomic data using publicly available single-cell data from localized prostate cancer.

**Supplementary Figure 7.** Loss of PTEN protein increases the transcriptional activity of the TGFβ response signature (TBRS) across different cell types (End-= Endothelial, T-= T-cells, Ma-= Macrophages).

## Supplementary Tables

**Supplementary Table 1.** Patient characteristics of the cohort, including age, Gleason grading, disease-free survival, BMI, and PTEN protein status and H-scores, among others.

**Supplementary Table 2.** Immunohistochemistry quality assessment results.

**Supplementary Table 3.** Gene Set Enrichment Analyses (GSEA) results for the human transcriptome comparing PTEN loss and PTEN presence.

**Supplementary Table 4.** Results of the Weighted Gene Co-expression Network Analyses (WGCNA) in the patient cohort.

**Supplementary Table 5.** Association results of different WGCNA modules identified in the patient transcriptomes with PTEN status, H-score, age, RNA quality (DV200), disease-free survival, purity, and PI3K-AKT-mTOR activity signature.

**Supplementary Table 6.** Results of differential expression analyses of the patient cohort.

**Supplementary Table 7.** Results of functional enrichment for the differentially expressed genes in the green module by gprofiler.

**Supplementary Table 8.** Results of *in silico* deconvolution by ESTIMATE.

**Supplementary Table 9.** Results of xCell *in silico* deconvolution of the human transcriptome dataset.

**Supplementary Table 10.** Calculation of differential estimates by xCell *in silico* deconvolution for PTEN loss and presence categories.

**Supplementary Table 11.** Results of the Weighted Gene Co-expression Network Analyses (WGCNA) in the mouse transcriptomes.

**Supplementary Table 12.** Association results of different WGCNA modules identified in the mouse transcriptomes.

**Supplementary Table 13.** Results of differential expression analyses of the mouse dataset.

**Supplementary Table 14.** Results of functional enrichment for the differentially expressed genes in the yellow mouse module by gprofiler.

