## Supplementary figures and images for "Transcriptional network analysis of PTEN protein-deficient prostate tumors reveals robust stromal reprogramming and signs of senescent paracrine communication"

### Supplementary Figure 3

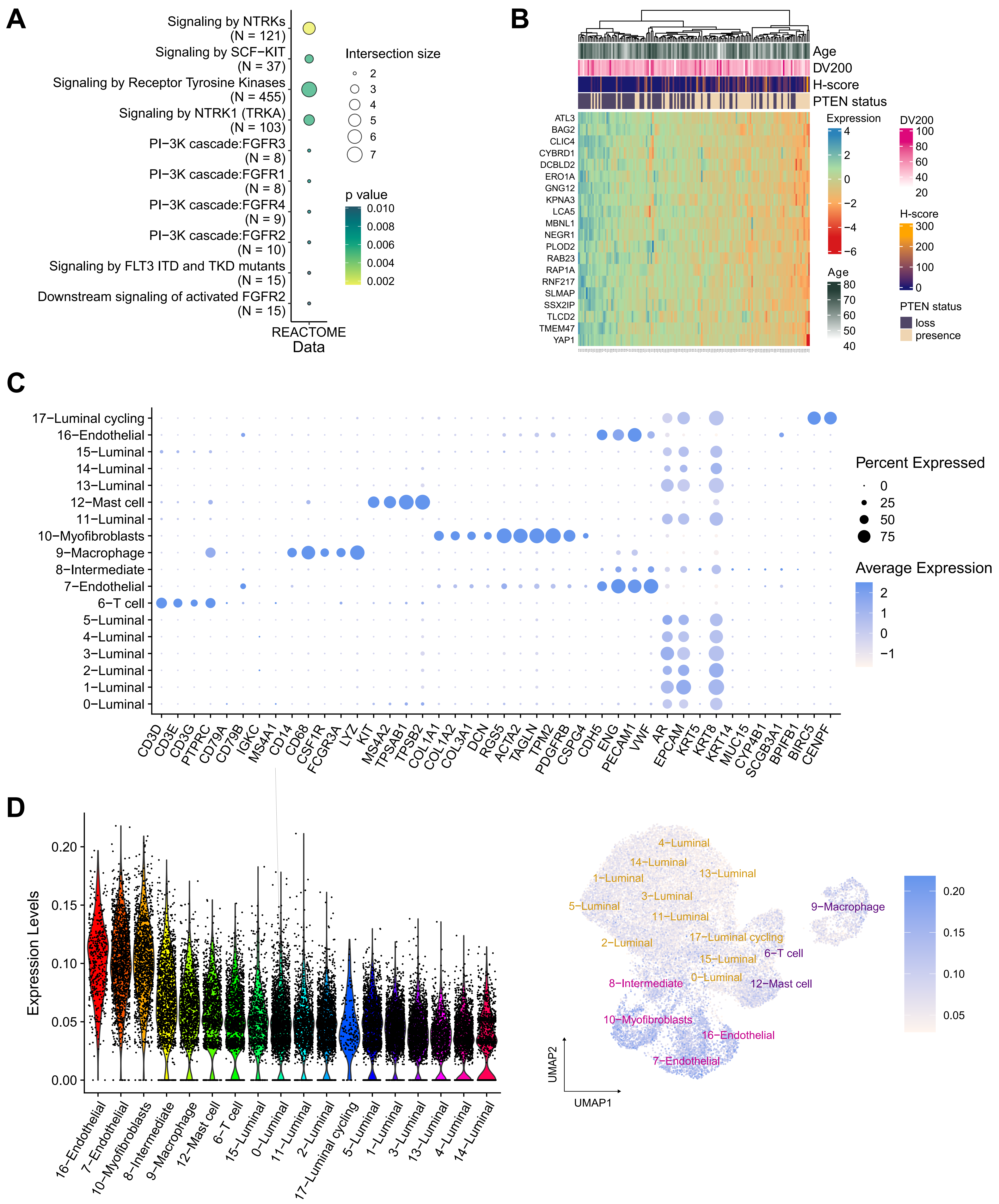

### Supplementary_Figure 1

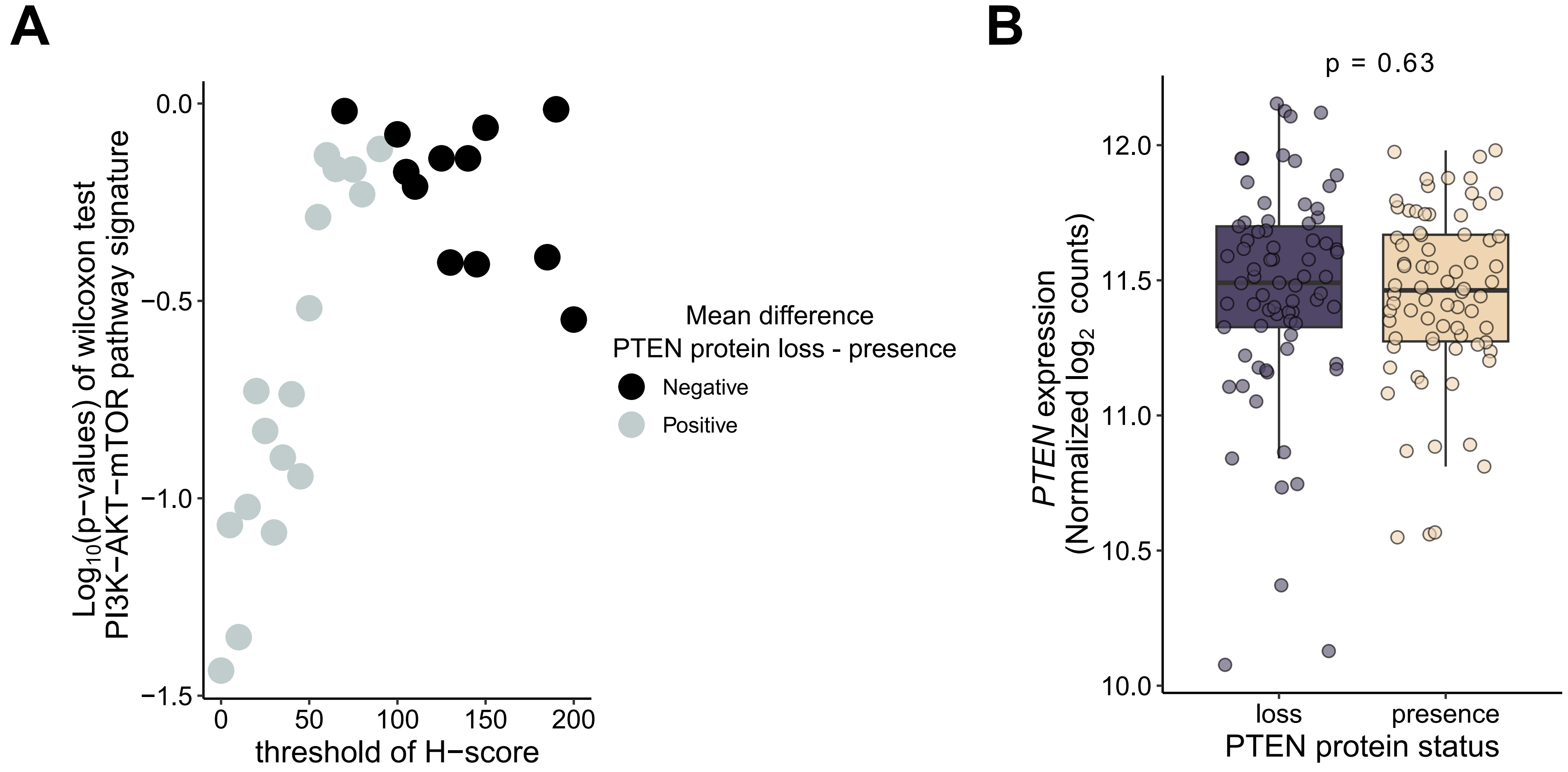

### Supplementary_Figure 2

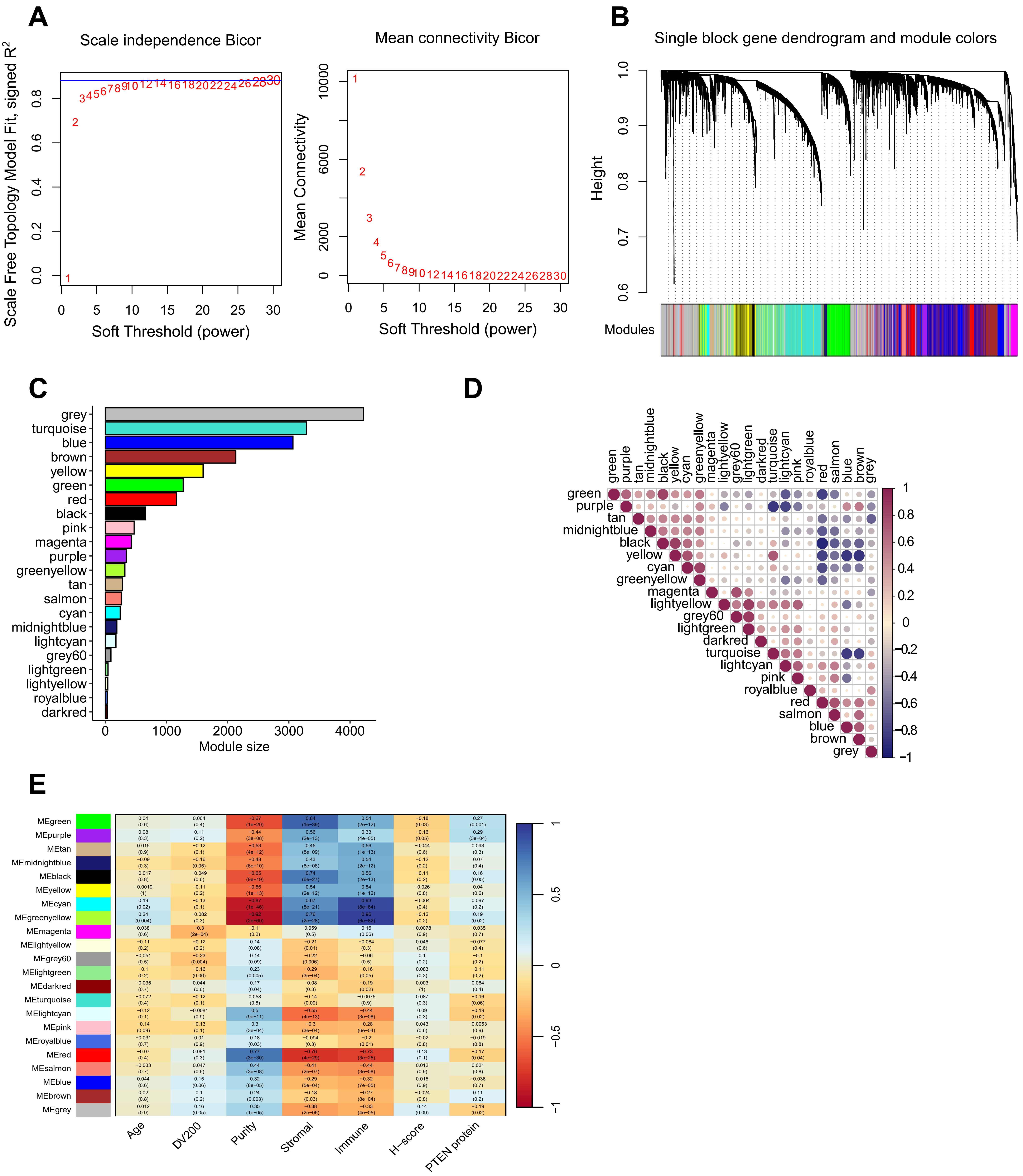

### Supplementary_Figure 4

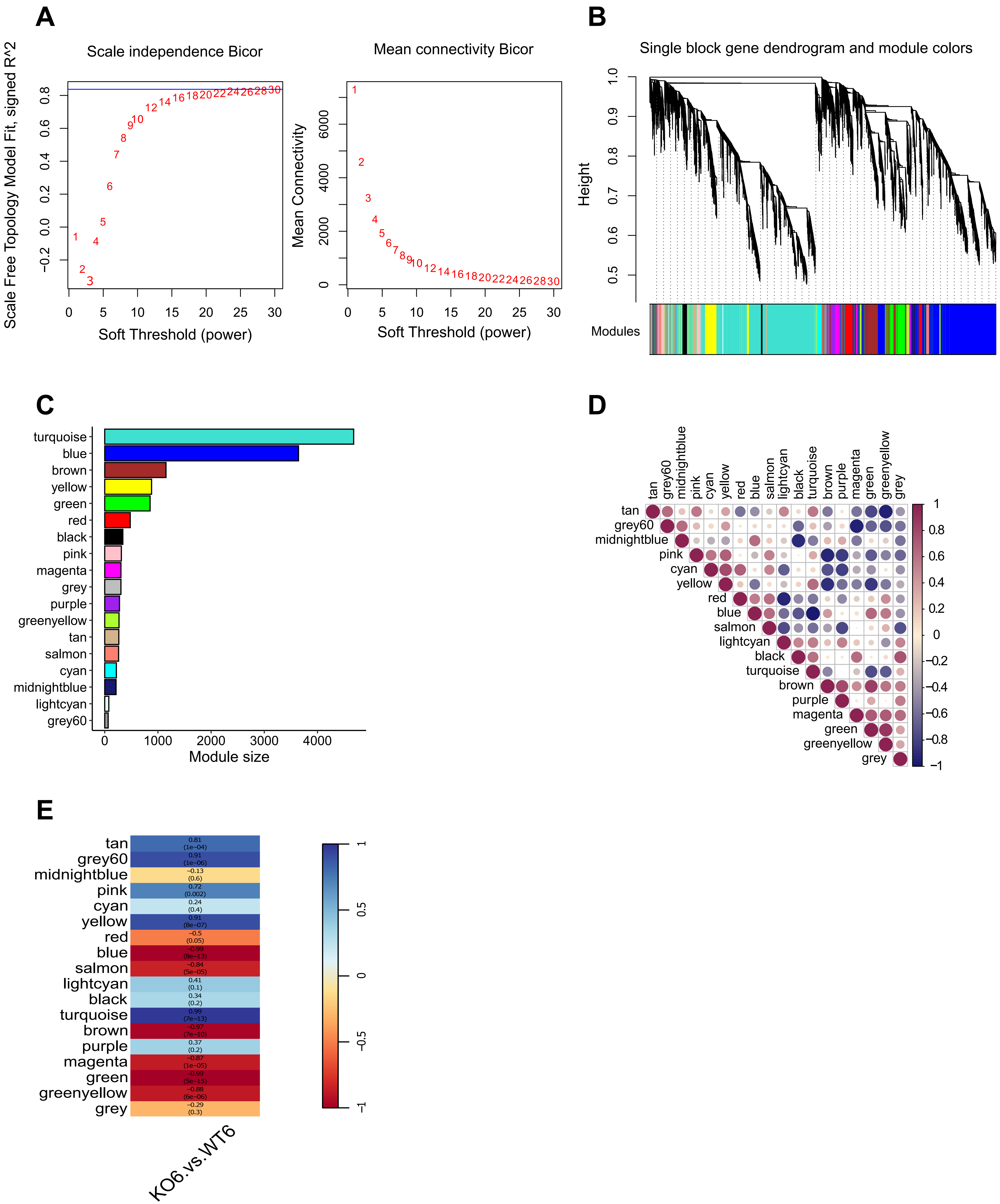

### Supplementary_Figure 5

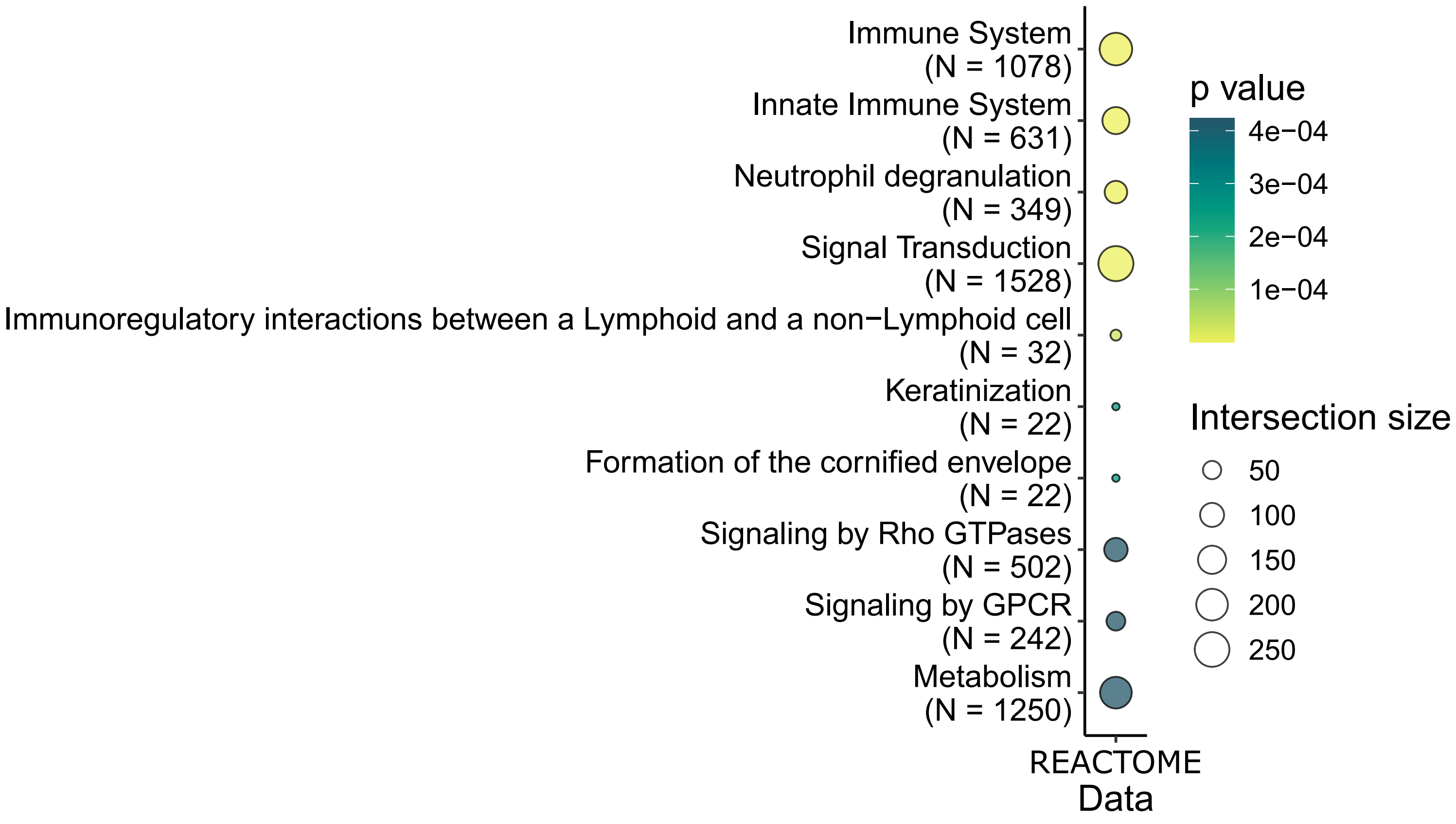

### Supplementary_Figure 6

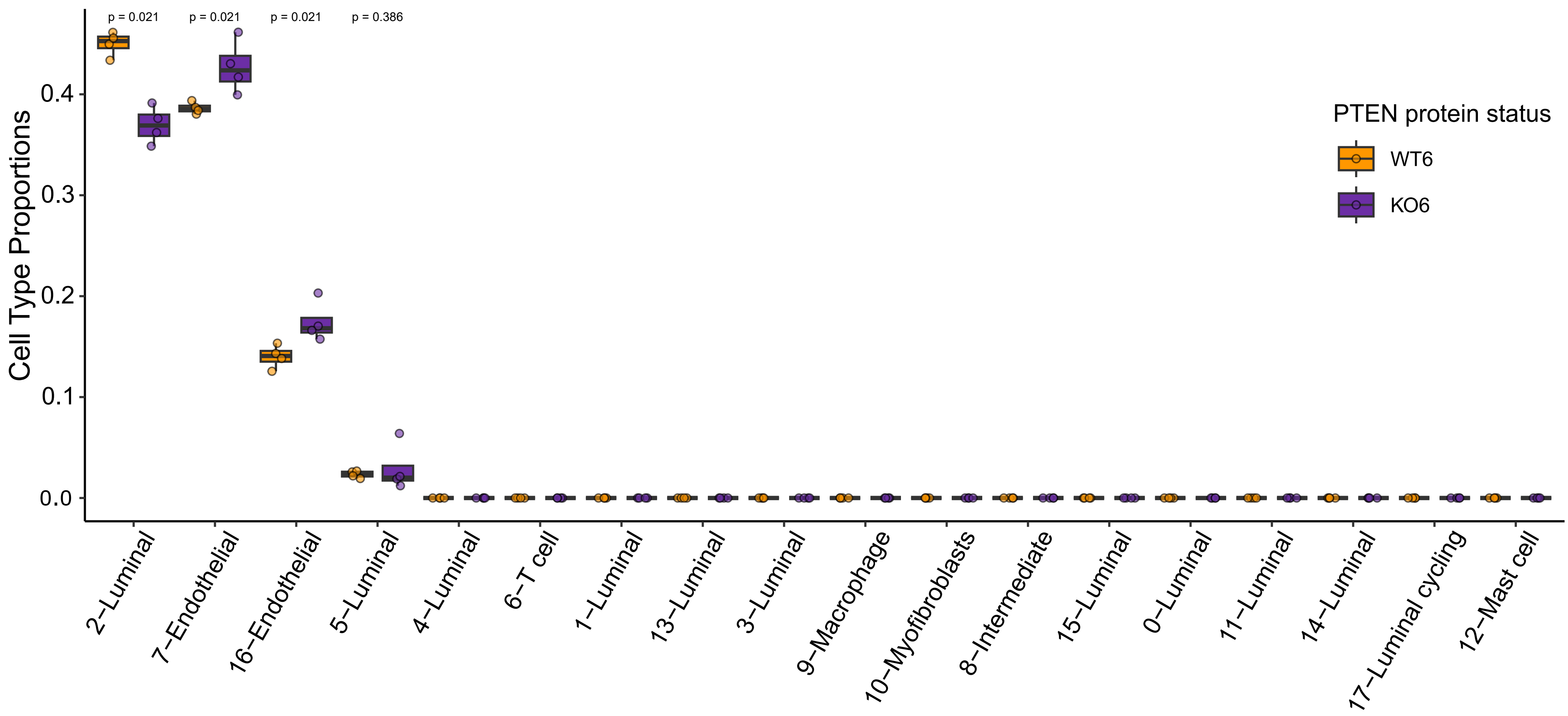

### Supplementary_Figure 7

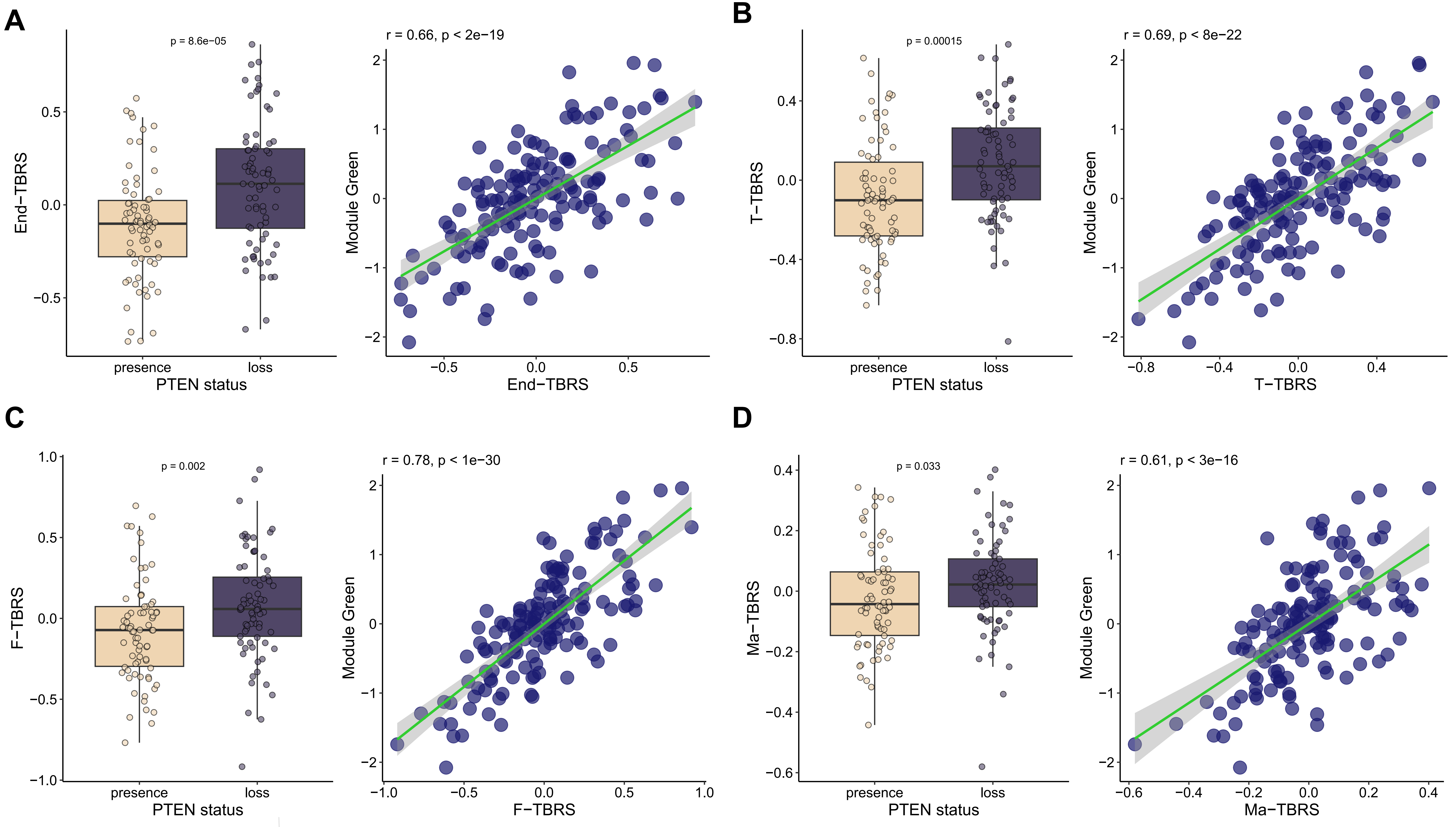
